# Audiomotor prediction errors drive speech adaptation even in the absence of overt movement

**DOI:** 10.1101/2024.08.13.607718

**Authors:** Benjamin Parrell, Minju Bae, Chris Naber, Olivia A. Kim, Caroline A. Niziolek, Samuel D. McDougle

## Abstract

Observed outcomes of our movements sometimes differ from our expectations. These sensory prediction errors recalibrate the brain’s internal models for motor control, reflected in alterations to subsequent movements that counteract these errors (motor adaptation). While leading theories suggest that all forms of motor adaptation are driven by learning from sensory prediction errors, dominant models of speech adaptation argue that adaptation results from integrating time-advanced copies of corrective feedback commands into feedforward motor programs. Here, we tested these competing theories of speech adaptation by inducing planned, but not executed, speech. Human speakers were prompted to speak a word and, on a subset of trials, were rapidly cued to withhold the prompted speech. On standard trials, speakers were exposed to real-time playback of their own speech with an auditory perturbation of the first formant to induce single-trial speech adaptation. Speakers experienced a similar sensory error on movement cancellation trials, hearing a perturbation applied to a recording of their speech from a previous trial at the time they would have spoken. Speakers adapted to auditory prediction errors in both contexts, altering the spectral content of spoken vowels to counteract formant perturbations even when no actual movement coincided with the perturbed feedback. Such adaptation was not observed when participants passively listened to perturbed feedback without the intention to speak, ruling out observational learning as the cause of adaptation in movement cancellation trials. These results build upon recent findings in reaching, and suggest that prediction errors, rather than corrective motor commands, drive audiomotor adaptation in speech.

## Introduction

Implicit motor adaptation is a prime example of the brain’s predictive capacities. In the standard framework, motor adaptation occurs when the predicted sensory consequences of a descending motor command are compared against feedback, and an internal model of those motor commands is adjusted to reduce observed discrepancies (i.e., sensory prediction errors) (Wolpert and Kawato, 1998; Wolpert and Flanagan, 2016; Krakauer et al., 2019). One lingering question in this framework is what serves as the input signal for the generation of sensory predictions that drive motor learning. In some models, that input signal is an efference copy generated from the descending motor command, which is then used to predict future states (Scott, 2004; Shadmehr and Krakauer, 2008; Houde and Nagarajan, 2011; Parrell et al., 2019). In other work, the input signal is defined as a higher-level motor plan (Day et al., 2016; Sheahan et al., 2016, 2018; McDougle et al., 2017; Vyas et al., 2020). It remains an open question whether motor planning or execution are sufficient for the computation of sensory predictions and errors, and whether there are common requirements for adaptation across motor subsystems.

In the case of speech adaptation, the prevailing idea is that ongoing feedback during actual execution of speech commands drives adaptation to audiomotor prediction errors by incorporating feedback-based motor commands into future movement plans (Tourville and Guenther, 2011; Guenther, 2016). Conversely, a recent model suggests that adaptation instead relies *directly* on auditory errors generated by the comparison of auditory reafference with auditory predictions deriving from a task or planning-level controller rather than by motor efference (Kim et al., 2023, 2025). To date, it has been difficult to distinguish between these competing theories as they make similar predictions in existing speech adaptation paradigms. That said, several lines of evidence from upper-limb adaptation support the notion that an “upstream” motor plan is sufficient to drive adaptation. One piece of evidence for this is the discovery that internal signals preceding a movement, such as planning “lead-in” and “follow-through” motions, can partition otherwise catastrophically interfering motor memories (Sheahan et al., 2016, 2018). Indeed, a range of similar contextual pre-cueing effects have been observed in motor adaptation, supporting the idea that preparatory signals seed predictive computations within the adaptation system (Wolpert et al., 2011; Howard et al., 2013; Heald et al., 2018, 2021; Vyas et al., 2020; Avraham et al., 2022; Churchland and Shenoy, 2024).

We have recently developed an experimental paradigm in limb control that is capable of teasing apart plan-vs. motor-based prediction error in sensorimotor adaptation, which we used to establish that implicit upper-limb motor adaptation could be induced in response to sensory errors that were unaccompanied by movements (Kim et al., 2022). Human subjects performed a “Go/NoGo” task that involved planning, but occasionally not executing, reaches to visual targets. On certain critical trials, subjects successfully inhibited reach commands but still observed a “virtual” sensory prediction error delivered via a visual cursor rotated relative to the planned reaching direction. These trials induced adaptation in subsequent movements, even though no movement had accompanied the error. In our view, these results offer particularly compelling support for the aforementioned plan-based model of adaptation.

Here, we applied this design to speech adaptation in order to answer one of the major outstanding questions about sensorimotor adaptation in speech: whether ongoing feedback from executed speech movements is necessary for adaptation or whether speech adaptation can leverage sensory predictions based only on planned speech acts. Based on our previous work in limb control, we expect to observe adaptation when speech motor movements are planned, but not executed, when paired with “virtual” auditory sensory errors. Such a result would strongly support models of speech adaptation driven directly by sensory prediction errors over models where adaptation is only driven through feedback-based motor commands. Moreover, by expanding this paradigm from a laboratory reaching task to speech, this work provides a critical test of the domain-generality of our previous findings. Speech differs in many ways from reaching, including sensory feedback modalities (auditory versus visual feedback), the neural basis of control (bulbar vs. spinal control) and the ultimate task goal (communication vs. interacting with the physical environment). If, despite these substantial differences, similar results are found in speech and reaching, this would strongly support a shared neurocomputational mechanism driving sensorimotor adaptation.

## Results

We implemented a task that allowed us to measure speech adaptation both with and without movement execution during perceived auditory prediction errors (Figure 2). Trials were organized into three-trial “triplets”, where the middle trial received a perturbation to the first vowel formant (F1), one of the resonances of the vocal tract determined by the position of the tongue, lips, and jaw which distinguish vowels from one another (Figure 2A). On “movement” trials, participants produced speech when cued by a visual stimulus. To measure adaptation in the absence of movement execution, we had participants occasionally inhibit a cued speech movement while still presenting them with a well-timed audiomotor prediction error (“no-movement” trials), using a formant perturbation applied to a recording of the speech produced on the immediately preceding trial. We measured adaptation for the movement and no-movement trials as the change in F1 from the trial preceding the perturbed trial to the two trials immediately following the perturbed trial (adaptation and aftereffects, Figure 2E).

**Figure 1.**
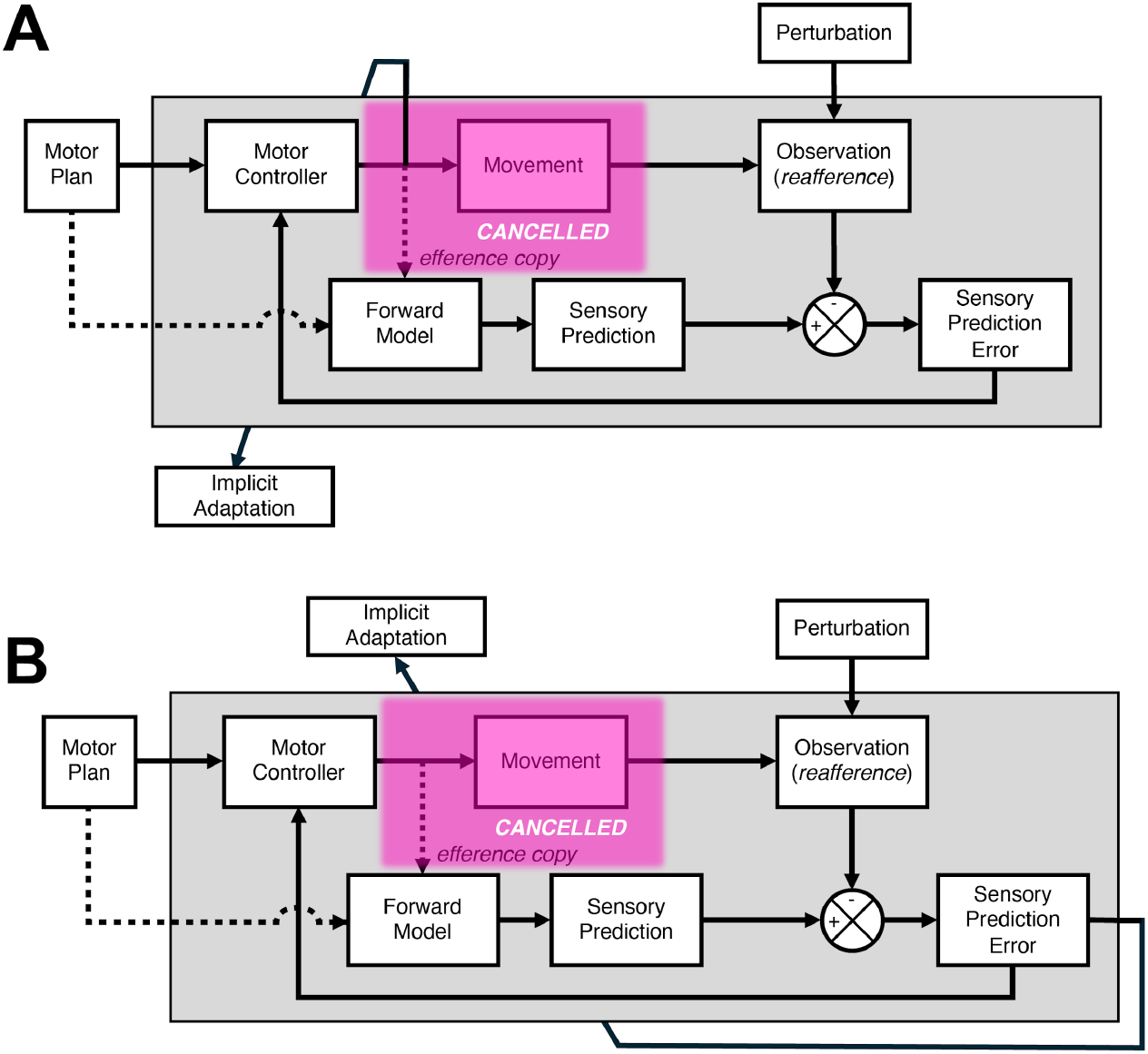
Schematic of competing theories of the source of sensorimotor adaptation. In both cases, reafferent feedback about movement outcomes is compared against sensory predictions generated by forward models, based either on an efference copy of motor commands or directly generated from the motor plan (dashed lines). Any discrepancies between observed and predicted sensory outcomes generates a sensory prediction error. **A**: In this model, sensory prediction errors drive adaptation indirectly: they are used by the motor controller to update motor commands (sensory feedback control), and these feedback-based motor commands are then used to adapt the motor control system. This is the basic mechanism of adaptation in the DIVA model of speech production (e.g., Guenther 2016). When movement is cancelled (pink box), this model predicts adaptation should not occur, as no feedback-based motor commands will be generated **B**: In this model, sensory prediction errors directly drive adaptation of the motor control system. As adaptation and feedback-based control are separated, this model predicts adaptation should occur even in the absence of motor movement as long as sensory prediction errors are still generated.

**Figure 2.**
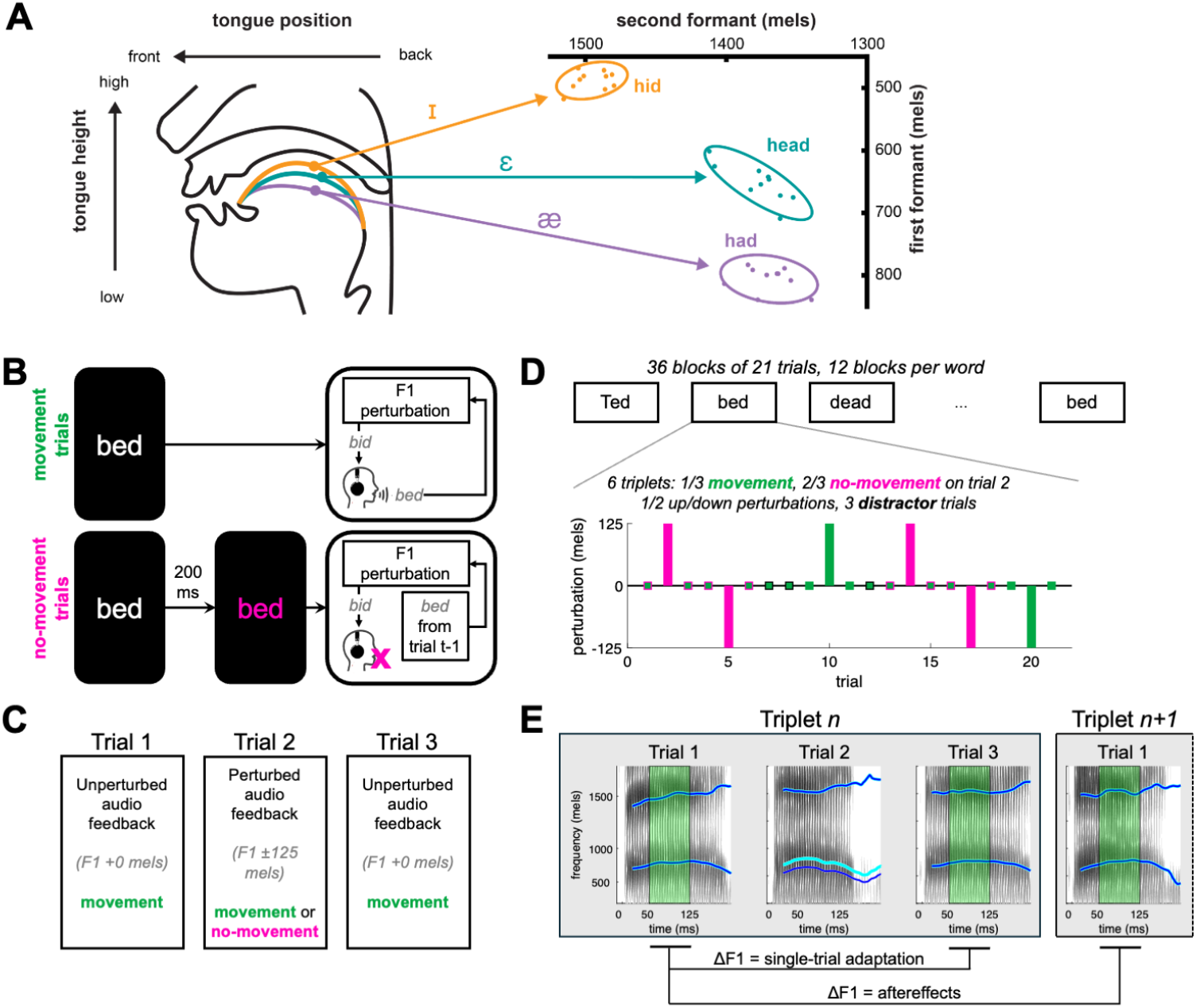
Experimental design. **A**: Illustration of the relationship between tongue position and vowel formants. The first two vowel formants (F1, F2) serve to distinguish between different vowels. **B**: Schematic of movement and no-movement trials. In movement trials, a stimulus word appeared on the screen and participants read the word out loud. Participants heard their own voice, with the first vowel formant (F1) perturbed or unperturbed. In no-movement trials, the stimulus word turned red 200 ms after appearing. Participants were instructed not to produce speech on these trials. Instead, the audio recording from the previous trial was played back to the participants with F1 perturbed. **C**: Triplet design. In each triplet, trials 1 and 3 were always movement trials with no perturbation applied. Trial 2 was variable – a movement or no-movement trial – and always had a perturbation applied. **D**: Experiment structure. The experiment consisted of 36 blocks which alternated between three stimulus words. Each block consisted of 6 triplets (2 with movement middle trials, 4 with no-movement middle trials) and 3 distractors (always movement trials) to disrupt the rhythm of the perturbations. **E**: Data analysis. Each trial shows a spectrogram of an example speech trial; cyan lines show produced vowel formants (F1 and F2) and blue lines show formants in headphone signal. For each trial, the produced F1 (the lowest frequency formant) was averaged between 50-125 ms after vowel onset (green shaded portion) to avoid coarticulatory effects of the initial consonant in each stimulus word and potential online compensatory effects. Single-trial adaptation was measured as the change in F1 from trial 1 to trial 3 within each triplet. Aftereffects were measured as the change from trial 1 of one triplet to the trial immediately following trial 3 of that triplet (either a distractor or trial 1 of the next triplet).

Despite the instruction to inhibit speech, participants nonetheless produced some acoustic speech material on a subset of no-movement trials (average of 22%, range 3-62%, Figure 2A). The prevalence of these inhibition failures suggests that our method was successful in driving participants to plan speech at the onset of the stimulus in these no-movement trials. In order to isolate learning in the absence of movement, only no-movement trials where participants successfully inhibited speech were used to measure adaptation.

Because the expected effect size of single-trial adaptation (∼3 mels in Hantzsch et al., 2022, Fig 3A) is much smaller than the expected standard deviation of F1 in typical speech (∼10-15 mels), our primary outcome measurement combined measurements taken from the trial immediately following the perturbation (adaptation) as well as the subsequent trial (aftereffects), in order to increase the power of our analysis and reduce noise. Participants produced robust single-trial learning following perturbations to both movement and no-movement trials (Figure 3A,B, main effect of perturbation direction F(1,10724) = 59.25, p < 1x10^-13^, partial R^2^ = 0.005). For movement trials, F1 following trials with a downward F1 perturbation was significantly higher (2.3 ± 0.8 mels) than following trials with an upward F1 perturbation (-6.2 ± 0.8 mels, t(10722) = 7.5, p < 0.0001, d = 0.23). Single-trial adaptation on movement trials was 3-4 times larger than in our previous work (Hantzsch et al., 2022), likely reflecting better control of stimuli in the current paradigm (see *Discussion*).

**Figure 3.**
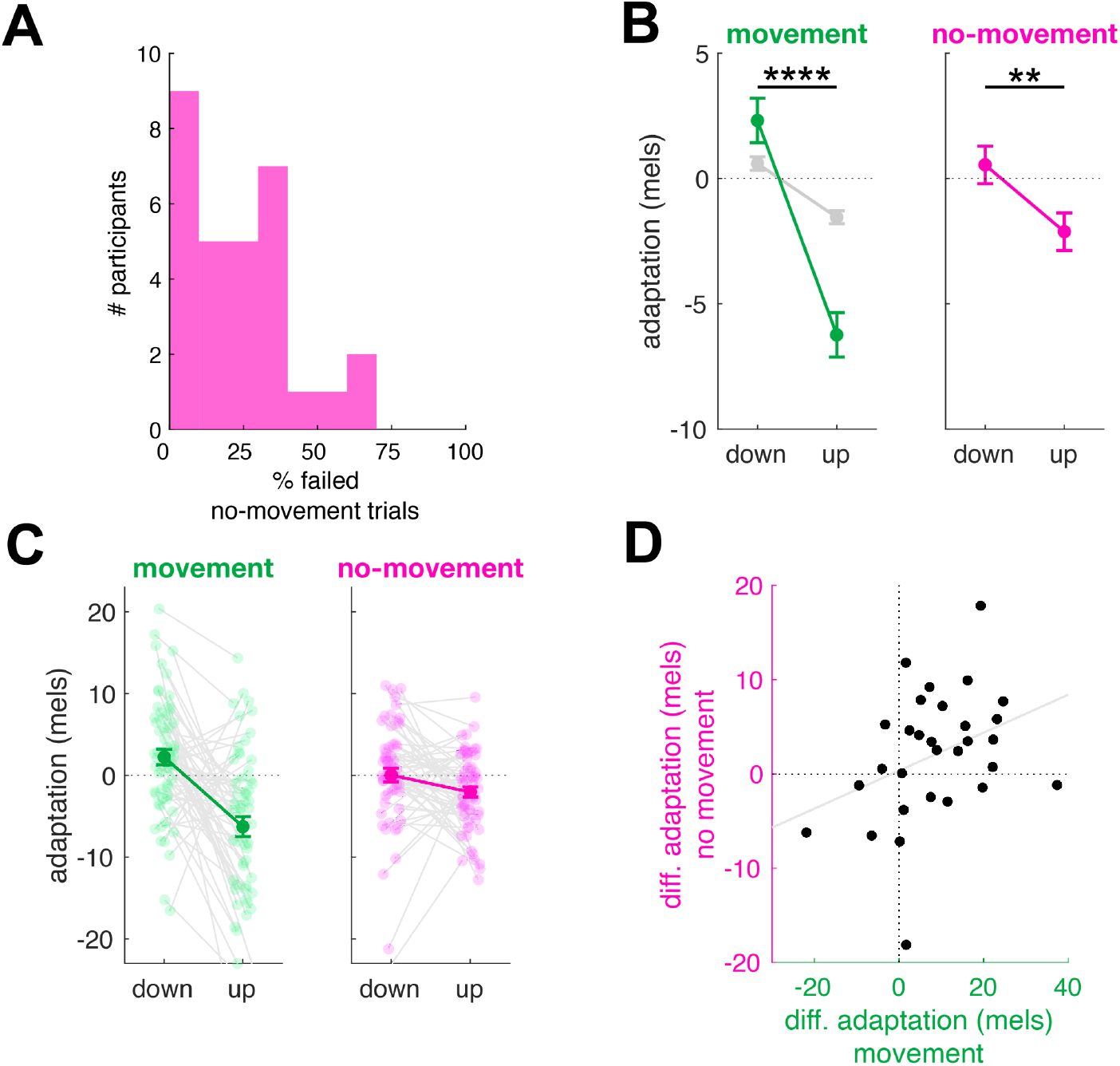
Experimental results for the adaptation index. **A**: The percentage of no-movement trials on which participants failed to inhibit vocal production. The average failure rate was 22% across participants. These trials were excluded from further analysis. **B**: F1 adaptation index (see Methods) following trials with downward and upward perturbations applied in movement (left) and no-movement (right) conditions. Estimated marginal means and standard errors shown in green and magenta. Gray indicates single-trial learning observed in Hantzsch et al. (2022). **C:** As for (B), showing data for individual participants. Means and standard error across participants are shown in dark, solid colors. Individual participant means shown as lighter colored dots connected by gray lines. **D**: Relationship across participants between the magnitude of differential adaptation (responses following downward perturbation minus responses following upward perturbations) in movement and no-movement conditions. ** p < 0.01, **** p < 0.0001.

Crucially, following no-movement trials, F1 was significantly higher in trials following downward perturbations (0.5 ± 0.7 mels) compared to upward perturbations (-2.1 ± 0.8 mels, t(10736) = 2.9, p = 0.004, d = 0.07). This result supported our hypothesis that speech adaptation could proceed without actual movement execution. The magnitude of adaptation was smaller in no-movement trials compared to movement trials, reflected by a significant interaction between movement condition and perturbation direction (F(1,10725) = 16.3414, p < 0.0001, partial R^2^ = 0.002). Overall, the magnitude of adaptation in no-movement trials was roughly one-third of that found in movement trials, similar to our previous results in adaptation of upper limb movements (Kim et al., 2022). (Potential reasons for this replicated difference in effect size are addressed in the *Discussion*.)

We then looked at a between-subjects correlation between adaptation effect sizes on movement versus no-movement triplets. Indeed, there was a moderate, though statistically marginal, positive relationship between learning in movement and no-movement trials across participants in the expected direction (Figure 3D, r = 0.36, p = 0.053).

As an additional analysis, we considered single-trial adaptation (in trials immediately following the perturbation, *t+1*) and aftereffects (in the next following trial, *t+2*) separately (Figure 4). The results for both adaptation and aftereffects were consistent with the combined results. For adaptation, there was a significant main effect of perturbation direction (F(1,5362.3) = 32.7, p < 1x10^-7^, partial R^2^ = 0.006) as well as a significant interaction between perturbation direction and movement condition (F(1,5362.3) = 10.2, p = 0.001, partial R^2^ = 0.002). The difference in F1 between upward and downward trials was significant for both movement (8.6±1.5 mels, t(5363) = 5.7, p < 0.0001, d = 0.24) and no-movement conditions (2.5±1.2 mels, t(5369) = 2.014 p = 0.04, d = 0.07), though the magnitude of this difference was again smaller for the no-movement condition.

**Figure 4.**
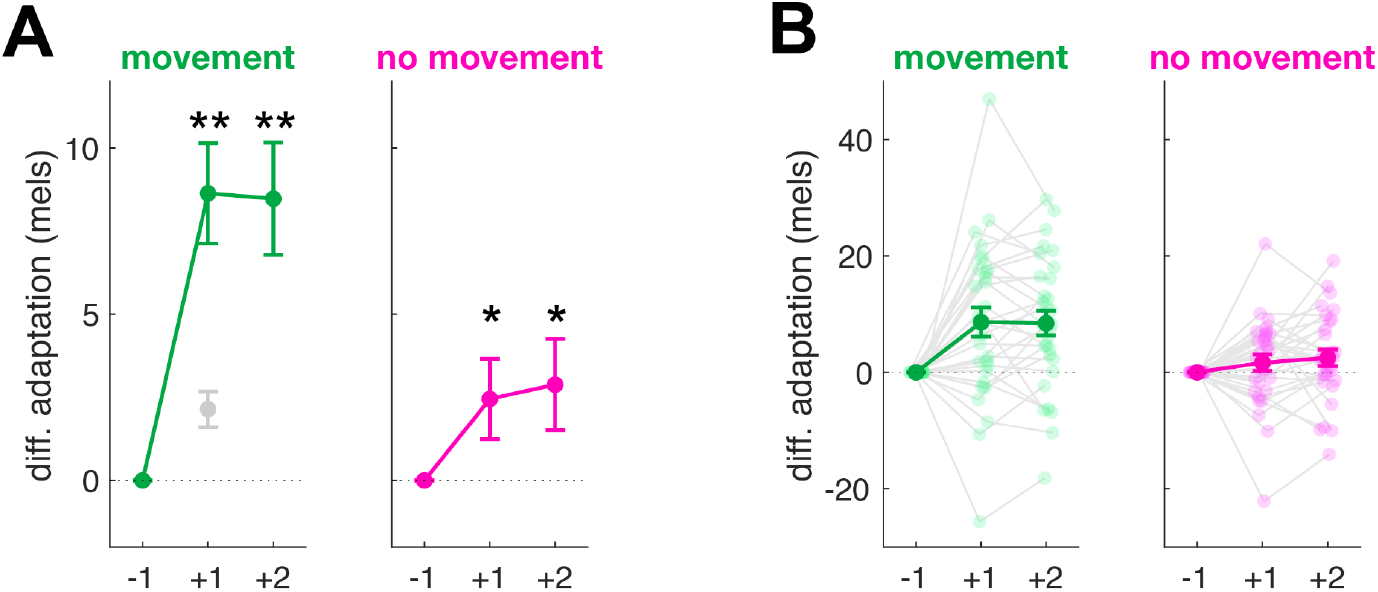
Experimental results separated by trial number. **A**: Differential adaptation shown separately for trials immediately following the perturbation (+1, adaptation) and the next trial (+2, aftereffects) in the movement (left) and no-movement (right) conditions. Estimated marginal means and standard errors shown in green and magenta. Gray indicates single-trial learning observed in Hantzsch et al. (2022). **B:** As for (A), showing data for individual participants. Means and standard error across participants are shown in dark, solid colors while individual participant means are shown as lighter colored dots connected by gray lines. * p < 0.05; ** p < 0.01.

The results for aftereffects were similar: Upward and downward perturbations led to differences in F1 in both movement (8.5±1.7 mels, t(5328) = 4.997 p < .0001, d = 0.22) and no-movement (2.9±1.4 mels, t(5341) = 2.110, p = 0.03, d = 0.07) conditions (main effect of perturbation direction: F(1,5355) = 27.2, p < 1x10^-6^, partial R^2^ = 0.005); and the magnitude of this difference was smaller in the no-movement condition (interaction: F(1,5355) = 6.6, p = 0.01, partial R^2^ = 0.001).

One possible explanation for the adaptation observed in no-movement trials is that participants might have learned from passive exposure to the auditory feedback error without necessarily generating a sensory prediction. In certain contexts, such observational learning has been observed in limb control (Mattar and Gribble, 2005; Pawlowsky et al., 2023). To test whether the adaptation observed in no-movement trials was a product of on observational learning we conducted a second study (Experiment 2) which mirrored the design of the initial study (Experiment 1), but substituted “observation” trials for no-movement trials. In this experiment, visual cues indicating whether to speak (movement trials) or remain silent (observation trials) were presented *prior* to stimulus onset, so that motor planning (and concurrent sensory prediction) would not occur in observation trials (Figure 5A).

**Figure 5.**
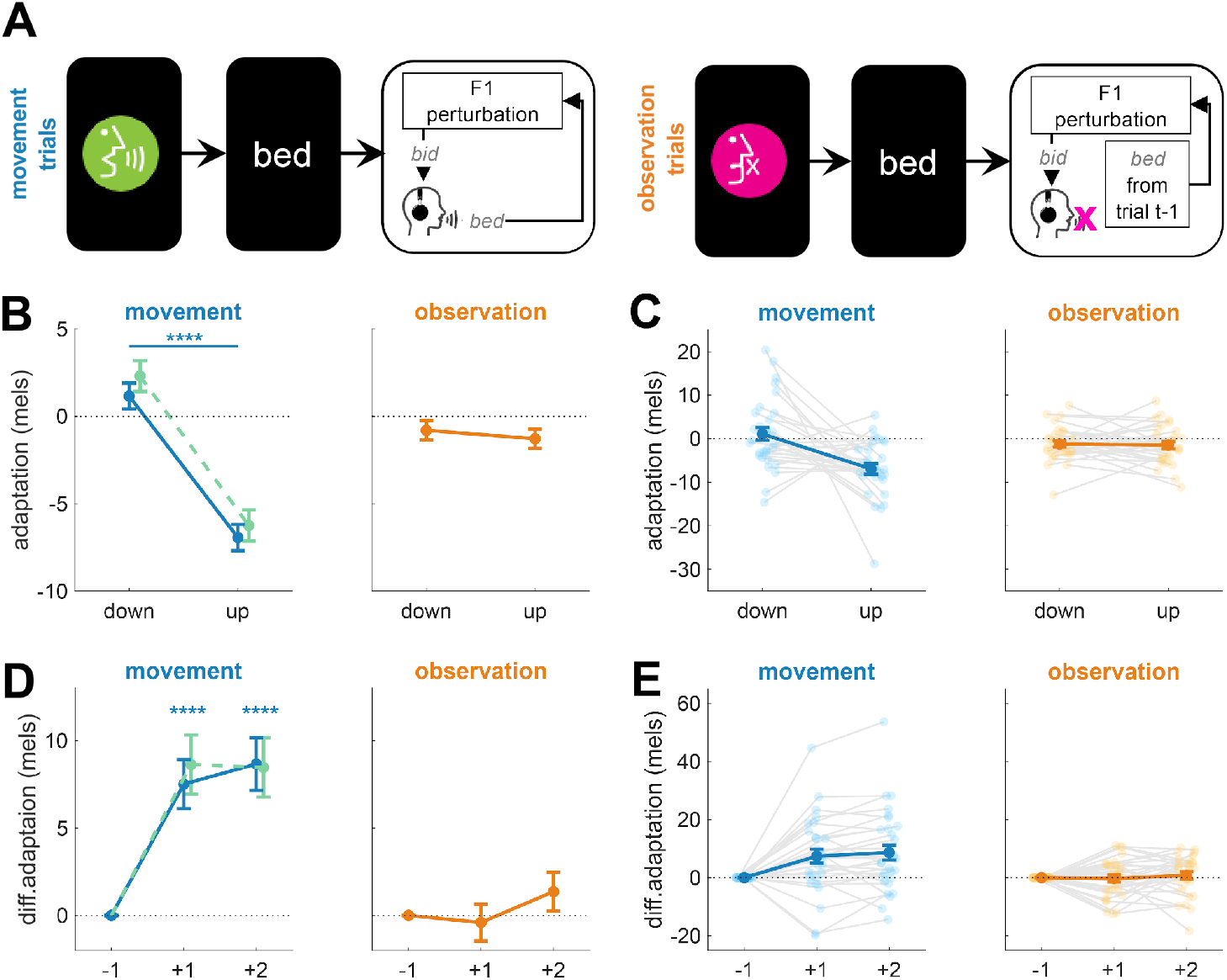
The experimental setup and results of Experiment 2. **A:** Schematic of movement (left) and observation (right) trials. In movement trials, a visual cue to speak appeared before a stimulus word, and participants read the word aloud. As in Experiment 1, participants heard their own voice with either perturbed or unperturbed F1 feedback. In observation trials, a cue to not speak was shown before a stimulus word, and participants heard the F1-perturbed audio recording from the previous trial. **B:** Estimated marginal means and standard errors of the F1 adaptation index (see Methods) following trials with downward and upward perturbations in the movement (left) and observation (right) conditions. Green represents the F1 adaptation index observed in movement trials in Experiment 1. **C**: As for (B), showing data for individual participants. Means and standard errors across participants are shown in dark, solid colors. Individual participant means shown as lighter colored dots connected by gray lines. **D:** Experimental results by trial number relative to the middle trial of each triplet. Estimated marginal means and standard errors of differential adaptation for the trial immediately following perturbation (+1, adaptation) and the subsequent trial (+2, aftereffects) in the movement (left) and observation (right) conditions. Green indicates corresponding results from Experiment 1 movement trials. **E**: As for (D), showing data for individual participants. Means and standard errors across participants are shown in dark, solid colors while individual participant means are shown as lighter colored dots connected by gray lines. **** p < 0.001.

We compared F1 changes following perturbations between movement and observation trials. Critically, adaptation was observed only in the movement condition (Figure 5B,C). In movement trials, F1 was significantly higher following downward perturbations (1.16 ± 0.75 mels) than following upward perturbations (−6.93 ± 0.75 mels; t(11824) = 7.84, p < .0001, d = 0.24). The effect seen here was a close replication of the movement condition in Experiment 1. In contrast, no significant difference was found in the observation condition (downward: −0.80 ± 0.56 mels; upward: −1.28 ± 0.55 mels; t(11846) = 0.63, p = .529). The contrast was reflected in a significant interaction between perturbation direction and movement condition (F(1,11842) = 35.27, p < .001, partial R^2^ = 0.003), suggesting that the adaptation observed in no-movement trials in Experiment 1 cannot be explained by observational learning alone.

Separate analyses of adaptation and aftereffects trials supported the same conclusion (Figure 5D,E). In movement trials, significant learning was observed in adaptation trials (t(5910) = 5.29, p < .0001, d = 0.23), as well as aftereffects trials (t(5882) = 5.79, p = < .0001, d = 0.256). However, in observation trials, neither adaptation (t(5933) = −0.38, p = .702) nor aftereffects (t(5907) = 1.24, p = .216) showed significant effects. The interaction between perturbation direction and condition was significant for adaptation (F(1,5919) = 20.09, p = .003, partial R^2^ = 0.003) and marginal for aftereffects (F(1,5909) = 15.47, p = .07, partial R^2^ = 0.003).

## Discussion

Here, we demonstrated that speech adaptation can occur even when audiomotor errors are not accompanied by movement execution. First, across two experiments we replicated the result that when participants were exposed to a perturbation of their first vowel formant on a single trial, changes in speech articulator movements that opposed that perturbation were visible on the two trials following the perturbation. These results confirm that single-trial speech adaptation observed in previous paradigms designed to examine online compensatory movements (Hantzsch et al., 2022) can be reliably elicited with our triplet design. Remarkably, adaptation occurred not only when participants heard an auditory perturbation during speech, but also when participants planned to produce speech but withheld overt speech movement and instead heard playback of their own speech from a previous trial with the perturbation applied. That is, when participants putatively generated a movement plan and a concomitant auditory prediction, they adapted their movements to correct for observed audiomotor prediction errors even though they did not produce overt speech movements in conjunction with those errors. Conversely, when motor planning was absent, as in observation trials, adaptation did not occur, ruling out observational learning as a mechanism for the observed changes in the no-movement trials. These results extend our recent findings in upper-limb visuomotor adaptation (Kim et al., 2022) to a new and very different motor domain: speech differs from reaching in sensory feedback modality (audition vs. vision), locus of neural control (bulbar vs. spinal), and movement goals (communication vs. environmental interaction). Despite these differences, the observed behavior in the two motor domains was strikingly similar. By dissociating observation from motor planning, the current study supports the idea that adaptation in speech, and likely more broadly across motor domains, involves a learning mechanism that depends on internal predictions generated during motor planning.

Why we failed to observe observational learning in Experiment 2, given that such learning has been seen in upper-limb motor control (Mattar and Gribble, 2005; Pawlowsky et al., 2023), remains an open question. We note that the precise conditions in the current study meaningfully differ from existing limb control studies on observational learning. For instance, a prior experiment that induced observational visuomotor reach adaptation (Mattar and Gribble, 2005) involved participant observation of the actual motor learning process, including corrective behaviors. In contrast, participants in the current study only observed altered sensory stimuli and were not allowed to observe an external learner. As another example, in Pawlosky et al. (2023), observational learning was seen when participants observed a sensory error (rotated cursor movement from a home position to a visually displayed target) while making an unrelated limb movement. While this is more similar to the current study in that no error-correction process was observed, it differs in whether or not participants were performing a motor action. Based on these distinctions, it may be that observing the learning process or engaging in an actual movement is critical for transferring observations to changes in action.

Alternatively, the difference in results between our speech adaptation data and others’ limb movement data may lie in the nature of the movement targets themselves. While limb control studies have well-defined motor targets (e.g., visually-displayed circles), speech targets are inherently individualized and flexible. In fact, speech has been shown to change by approximating the acoustic characteristics of perceived speech produced by other talkers (Goldinger, 1998; Shockley et al., 2004; Babel, 2012; Babel and Bulatov, 2012). In the current experiment, it is unclear whether participants perceived the perturbed auditory feedback as an external speaker’s voice, which could drive phonetic convergence even in passive social situations (Goldinger, 1998; Namy et al., 2002). If the altered feedback was perceived as another talker, it is possible that phonetic convergence may have been elicited in our study, thereby counteracting any observation-based adjustment and resulting in the non-significant net adaptation (though the reduction in convergence with a delay between the auditory stimulus and vocal productions seen in previous work (Goldinger, 1998) may make this explanation unlikely). In any case, the striking difference in behavior seen in no-movement trials, which elicit clear adaptation, and observation trials, which do not, strongly suggests that motor planning enhances audiomotor adaptation in speech, presumably by supporting the generation of a sensory prediction error, which is likely not present in observation trials.

The current results suggest that audiomotor speech adaptation is driven, at least in part, by using sensory prediction errors to directly update internal models controlling speech articulators. Such updates are consistent with dominant models of sensorimotor learning in limb and oculomotor control (Wolpert et al., 1998; Smith et al., 2006; Shadmehr and Krakauer, 2008; Shadmehr et al., 2010; Hadjiosif et al., 2021) and with a recent model of speech motor learning (Kim et al., 2023). However, these results are inconsistent with the dominant model of sensorimotor adaptation in speech production, which suggests that adaptation is driven by the incorporation of corrective movements generated by a feedback controller in response to sensory errors into future feedforward motor programs (Tourville and Guenther, 2011; Guenther, 2016; Kearney et al., 2020), which itself is partially based on other ideas in limb control (Kawato et al., 1987; Albert and Shadmehr, 2016).

Notably, the magnitude of the change observed on the movement triplets (∼8 mel difference between upward and downward perturbations) was roughly 2-3 times larger than we previously observed using the same analysis window (Hantzsch et al., 2022). Why did we see this large increase in effect size? One possibility is that our new design, which uses blocked repetition of the same stimuli, induces stronger learning signals than when stimuli are mixed across trials as in previous studies. This could be because adaptation in speech only partially generalizes to untrained words (Rochet-Capellan et al., 2012; Caudrelier et al., 2018), similar to the local spatial generalization of learning observed in reaching (Gandolfo et al., 1996; Krakauer et al., 2000; Donchin et al., 2003). Importantly, the larger effect size we observed in our movement trials may allow for more precise assays of factors that affect sensorimotor adaptation in speech, given that around 100 observations of learning can be obtained in the time it would typically take to obtain a single observation (e.g., mean asymptotic learning) using a more traditional paradigm with extended exposure to a repeated auditory perturbation.

Intriguingly, the size of the reduction we observed in speech adaptation between movement and no-movement trials (roughly one-third, Figures 2 and 3) was very similar to the reduction previously observed in a similar reaching task (Kim et al., 2022), pointing to a domain-general explanation. We see several possible explanations for this observed reduction: First, it has been established, both in speech and reaching tasks, that temporal delays between movement and sensory feedback substantially impair adaptation to errors (Kitazawa et al., 1995; Brudner et al., 2016; Schween and Hegele, 2017; Zhou et al., 2017; McDougle and Taylor, 2019). In speech, the magnitude of adaptation is reduced by roughly 50% when auditory feedback is delayed by only 100ms, and adaptation is essentially absent when delays reach 250-500ms (Max and Maffett, 2015; Shiller et al., 2020). This suggests that the critical comparison driving auditory error processing is highly temporally-specific, particularly in speech. In our paradigm it is impossible to know exactly when speech would have occurred on the no-movement trials; as an estimate, the latency of the speech feedback on a given no-movement trial was matched to the latency on a previous movement trial. This method is likely to have introduced variance between the timing of anticipated and perceived auditory feedback on no-movement trials, potentially leading to a reduction in the magnitude of adaptation.

Second, reductions in somatosensory feedback inherent in no-movement trials may have suppressed the degree of adaptation possible. It has been suggested that, in limb control, adaptation may result not from a drive to reduce visual sensory error but rather from the recalibration of somatosensory and other sensory signals (Tsay et al., 2022). Consistent with this idea, putatively disruptive noninvasive stimulation to primary somatosensory cortex following visuomotor reach adaptation substantially decreases how much of this learning is retained (Ebrahimi and Ostry, 2024). Thus, when errors occur in the absence of somatosensory feedback, such as in our no-movement trials, it is possible that adaptation magnitude would be reduced.

Finally, it may also be the case that adaptation in speech can be driven by auditory prediction errors both directly through updates to predictive internal models and indirectly through incorporation of previous feedback-based commands into feedforward motor programs. In fact, there is some support for this “dual input” idea in tasks requiring adapting to dynamic force-field perturbations during reaching (Albert and Shadmehr, 2016). Thus, an important note here is that while many results point to a plan-based sensory prediction error model of adaptation, both planning and ongoing feedback commands could both provide inputs that generate sensory predictions. Future work can directly assay if and how both sources of information fuel adaptation.

Although direct neurophysiological data related to speech adaptation is limited, our results are consistent with invasive and non-invasive imaging studies that have examined sensory prediction in speech. When we speak, auditory cortical activity is suppressed relative to when we passively listen to the same sounds, a process thought to be driven by the cancellation of auditory reafference using predictive internal models (Curio et al., 2000; Houde et al., 2002; Ventura et al., 2009; Flinker et al., 2010). This suppression effect is reduced in less prototypical productions, suggesting that the prediction is based on a plan (or target) rather than motor efference (Niziolek et al., 2013; Beach et al., 2024; Tang et al., 2025). Recent work has shown that suppression is modulated during adaptation, and that the degree of modulation predicts the degree of learning, strongly suggesting that these predictions (and the resulting sensory prediction errors) play a critical role in adaptation (Kim et al., 2025). Single-unit recordings in marmosets indicate that suppression of auditory cortex firing begins hundreds of milliseconds ahead of vocalization initiation (Eliades and Wang, 2003), and stimulus-specific cortical modulation has been demonstrated in this pre-vocalization window in humans (Daliri and Max, 2016), suggesting that vocalization planning alone can modulate auditory cortex activity and may be sufficient to support error computation.

Although we did not directly compare the magnitude of adaptation in response to the upwards and downwards perturbations, there is a clear asymmetry visible in the results, with responses to upward perturbations being much larger than responses to downward perturbations. The source of this asymmetry is not immediately apparent. However, similar asymmetries have been observed before, including in our previous study of single-trial adaptation across a large sample of speakers, though the direction of the asymmetry varies across studies and across vowels (Mitsuya et al., 2015; Niziolek and Parrell, 2021; Hantzsch et al., 2022; Zeng et al., 2025). Notably, by focusing on the difference between adaptation to opposing formant perturbations, we were able to reliably detect learning despite this asymmetry in response magnitude.

There are two caveats about our no-movement condition. First, although single-trial adaptation was robust in the no-movement condition, the overall effect was relatively small (∼2 mels, d = 0.07). Though this magnitude is roughly equivalent to our previous work demonstrating single-trial adaptation (in movement trials) when stimulus words varied across trials (Hantzsch et al., 2022), speech, in general, is substantially more variable than this, with a standard deviation in F1 of roughly 10-15 mels. Our triplet design, however, allowed for a high number of observations per participant in order to look for this modest effect. Second, while we excluded no-movement trials with any overt sound production of any kind, it is possible that participants nonetheless produced some very subtle muscular activity on a subset of these trials. However, there is some evidence, though from a slightly different task, that articulatory movements and voicing frequently co-occur when stop signals occur at a similar latency after a go signal as in the current study (Tilsen and Goldstein, 2012). This suggests that the acoustic signal used as a criteria for exclusion here is likely to also exclude trials with overt movement – as such, we do not believe that potential latent muscular activity substantially changes the main conclusions we draw from these data. Nonetheless, future work could measure muscle activity or movement more directly, such as with surface EMG of the masseter or articulatory tracking of the tongue, to address this question more directly.

Overall, our study suggests that the direct updating of internal models through sensory prediction errors is sufficient to drive speech adaptation. In our view, speech planning may provide the critical sensory predictions that, when violated, lead to adaptation of internal models governing speech control. These results do not support models of speech adaptation that rely solely on the incorporation of feedback-based motor commands into future feedforward plans, suggesting that such models could be revised to incorporate direct updating of internal models through sensory error processing. Moreover, by extending our previous findings in upper-limb adaptation (Kim et al., 2022) to a novel motor domain (speech) and a different sensory system (audition), we show that plan-based predictions may form the basis for sensorimotor adaptation across a wide range of human motor behaviors, pointing to a shared, domain-general neurocomputational mechanism.

## Methods

### Participants and power analyses

30 participants were tested in each experiment (Experiment 1: 5 male/25 female, age range 18-45, mean age 23.4; Experiment 2: 4 male/26 female, age range 18-64, mean age 25.6). The sample size was determined using a bootstrapping procedure with effect sizes observed in our previous work (Hantzsch et al., 2022), with the target of 90% power to detect a similar sized effect at α = 0.05. All participants were native speakers of American English, without any reported history of neurological, speech, or hearing disorders. All participants passed an automated Hughson-Westlake hearing screening (thresholds 25 dB HL or lower in both ears at 250, 500, 1000, 2000, and 4000 Hz). Participants gave informed consent prior to participation in the study and were compensated either monetarily or with course credit. All procedures were approved by the Institutional Review Board of the University of Wisconsin–Madison.

### Task setup

Participants were seated in a sound-insulated booth in front of a computer monitor. On each trial, a target word appeared on the screen in white text (Figure 1B). Each trial lasted 1.7 s from stimulus onset. Trials were separated by 1.25 s plus a random delay of 0-0.5 s. Participants were instructed to read the words as quickly as possible as they appeared on each trial. Two trial types were used – “movement” trials and “no-movement” trials, following our previous study in reach adaptation (Kim et al., 2022). On the majority of trials (movement trials), the word stayed white for the duration of the trial. On a subset of trials (no-movement trials), the target word turned red 200 ms after it appeared and stayed red for the remainder of the trial. Participants were instructed to not produce overt speech if and when the target word turned red.

200 ms was chosen after pilot testing suggested that this delay results in the inhibition of speech on most, but not all, trials in the majority of participants. This delay was therefore long enough to elicit movement planning but short enough to enable mostly successful inhibition, thereby allowing us to test whether sensory feedback given in the absence of overt movement could drive speech adaptation.

On movement trials, participants’ speech was recorded (AKG C520), digitized with a USB sound card (Focusrite Scarlett 2i2), processed through the Audapter software package (Cai et al., 2008; Tourville et al., 2013), and played back to the participants over closed-back, over-the-ear headphones (Beyerdynamic DT 770). Speech was played back at a volume of approximately 83 dB SPL and mixed with speech-shaped noise at approximately 60 dB SPL. The final level of the playback speech signal varied with the amplitude of participants’ produced speech. The noise, combined with the closed-back headphones, served to minimize potential perception of the participants’ own unaltered speech, which may have otherwise been perceptible through air or bone conduction. The latency of audio playback on our system is ∼18 ms, as measured using the protocol suggested by (Kim et al., 2020).

Trials were organized into “triplets.” The first and last trials of each triplet were always movement trials with veridical auditory feedback (Figure 1C). On the middle trial of each triplet, the auditory feedback participants received was always perturbed, such that the first vowel formant (F1, Figure 1A) was either raised or lowered by 125 mels (a perceptually-calibrated measurement of frequency) throughout the trial using Audapter. Middle trials were either movement trials (? of triplets) or no-movement trials with a stop signal (⅔ of triplets). On perturbed movement trials, the auditory feedback was perturbed in real time by identifying the vowel formants using linear predictive coding (LPC) and filtering the speech signal to introduce a shift to those formants. On perturbed no-movement trials, Audapter was used offline to apply the same shift to a recording of the participants’ production from the previous trial (i.e. the first trial of the triplet, which was always an unperturbed “movement” trial). On these trials, Audapter was initiated at the same time as on movement trials, but played back this perturbed speech signal rather than playing back the speech recorded in real time. Thus, the latency of the auditory feedback on each no-movement trial was the same as on the immediately preceding trial.

Triplets were organized into blocks (Figure 1D). Each block contained 2 movement triplets and 4 no-movement triplets. Upward and downward frequency perturbations were balanced within each block. The order of triplets within each block was randomized. Each block additionally contained 3 distractor trials, randomly inserted between triplets, in order to disrupt the rhythm of the perturbations across trials. All distractor trials were movement trials with veridical auditory feedback. Each block used a single stimulus word (“bed”, “dead”, or “Ted”, all sharing the same target vowel /ε/); stimuli were pseudo-randomized across blocks such that there were an equal number of blocks for all stimuli, and no consecutive blocks used the same stimulus. The experiment consisted of 36 total blocks, yielding 216 total triplets. This resulted in 36 perturbations in movement blocks and 72 perturbations in no-movement blocks for each perturbation direction. A short self-timed break was allowed between each block.

In order to encourage participants to produce their speech movements quickly, they were given points based on their response latency on movement trials. 10 points were given for responses with latencies up to 500 ms, falling off by 1 point every 40 ms thereafter. To encourage participants to inhibit overt speech production on no-movement trials, -25 points were given when a spoken response was detected. On all trials, the onset of speech (and response latency) was determined as the point where the amplitude of the microphone signal crossed above a predetermined, low amplitude threshold. Prior to the experiment, participants were trained in this paradigm: after a verbal description of the task, participants completed a single block of trials (21 total trials). This training was repeated as needed until participants showed the ability to respond appropriately on the no-movement trials (training repeated if participants produced any vocalization on more than 2/4 no-movement trials).

A second experiment (Experiment 2) followed the same design, with “observation” trials used instead of “no-movement” trials (Figure 4A). In this experiment, either green or pink visual cues were presented before stimulus words. The green picture of someone speaking indicates ‘speak’, and participants were instructed to read the words aloud, while the pink picture of someone with their mouth closed indicates ‘don’t speak’, and participants were instructed to remain still. Since these cues were presented before each stimulus word, observation trials would not elicit movement planning, and participants instead passively listened to a recording of their own speech from the previous trial with the formant perturbation applied, as in “no-movement” trials in Experiment 1.

### Data quantification

Formant data were tracked using wave_viewer (Niziolek and Houde, 2015), which provides a MATLAB GUI interface for formant tracking using Praat (Boersma and Weenink, 2019). LPC order and pre-emphasis values were set individually for each participant. Vowels were initially automatically identified by locating the samples which were above a participant-specific amplitude level. Subsequently, all trials were hand-checked for errors. Errors in formant tracking, identified by discontinuities in the formant tracks or values outside a vowel’s typical range, were corrected by adjusting the pre-emphasis value or LPC order. Errors in the location of vowel onset and offset were corrected by hand-marking these times using landmarks in the audio waveform and spectrogram. For each trial, F1 was averaged from 50-125 ms after vowel onset (Figure 1E) to avoid 1) coarticulatory influences on vowel formants from the initial consonant and 2) potential changes in vowel formants due to online feedback corrections, which begin roughly 150 ms after vowel onset (Cai et al., 2012; Parrell et al., 2017). Recent work has shown that this window is the most likely to accurately capture learning in this single-exposure paradigm (Hantzsch et al., 2022).

Adaptation was quantified in three ways. Single-trial adaptation was measured as the change in F1 (in mels) from the first to the third trial of a triplet (i.e., after exposure to the auditory perturbation). We additionally calculated single-trial aftereffects – the retention of learning – by measuring the change in F1 (in mels) from the first trial in a triplet to the trial immediately following the third trial of that triplet. Finally, we computed an “adaptation index,” which was quantified by simply combining the above two metrics for each triplet. In all cases, the critical trials were unperturbed movement trials, either the first trial of the following triplet or a distractor trial. Because of the rather high variance in F1 production for vowels (standard deviation of ∼15-20 Hz, Whalen and Chen, 2019) relative to the maximum expected effect size given previous work (∼2 Hz, Hantzsch et al., 2022), the adaptation index was our primary dependent variable of interest, designed to maximize our statistical power and potential to detect a small effect relative to the expected variance in production.

A small number of trials were excluded due to errors in production (i.e., the participant said the wrong word), disfluencies, or unresolvable errors in formant tracking (Experiment 1: 0-3.6% across participants, median .03%, Experiment 2: 0-6%, median .07%). Additionally, no-movement triplets or observation triplets where participants produced any detectable vocalization on the middle perturbed trial were excluded in order to isolate learning effects when no speech movement was produced (Experiment 1: 4-89 triplets excluded across participants, median 32, Experiment 2: 0-115 triplets excluded, median 3). In total, these exclusions resulted in 64-72 independent measurements of movement triplets (median 71) and 52-140 independent measurements of no-movement triplets for each participant (median 111) in Experiment 1 and 63-72 independent measurements of movement triplets (median 70) and 27-140 independent measurements of observation triplets (median 137) in Experiment 2.

## Statistical analysis

Following our previous study in reaching (Kim et al., 2022), linear mixed effects models were run using the lme4 package in R (Bates et al., 2015). For single-trial adaptation, retention, and the adaptation index models, model predictors included perturbation direction (upward or downward), triplet type (movement or no-movement), and the interaction between these two factors. All trials were included in the model, with Participant was included as a random effect (intercept) to control for multiple observations from each individual participant. Statistical significance was assessed using the lmerTest package (Kuznetsova et al., 2017). Post-hoc comparisons were conducted with estimated marginal means using the Satterthwaite method for approximating the degrees of freedom (Lenth et al., 2023). Effect sizes are reported as partial R^2^ values from the linear mixed effects models and Cohen’s *d* for pairwise comparisons. Summary statistics report means and standard errors.

## Acknowledgments

Work supported by grants R01 NS134754 (C.A.N, B.P, & S.D.M.), R01 DC019134 (C.A.N. & B.P.), R01 DC017091 (B.P.), and R01 NS132926 (S.D.M.) from the National Institutes of Health and BCS 2120506 (C.A.N. & B.P.) from the National Science Foundation.

